# Long-term persistence of crAss-like phage crAss001 is associated with phase variation in *Bacteroides intestinalis*

**DOI:** 10.1101/2020.12.02.408625

**Authors:** Andrey N. Shkoporov, Ekaterina V. Khokhlova, Niamh Stephens, Cara Hueston, Samuel Seymour, Andrew J. Hryckowian, Dimitri Scholz, R. Paul Ross, Colin Hill

## Abstract

The crAss-like phages are ubiquitous and highly abundant members of the human gut virome that infect commensal bacteria of the order Bacteroidales. Although incapable of classical lysogeny, these viruses demonstrate unexplained long-term persistence in the human gut microbiome, dominating the virome in some individuals. Here we demonstrate that rapid phase variation of alternate capsular polysaccharides plays an important role in dynamic equilibrium between phage sensitivity and resistance in *B. intestinalis* cultures, allowing phage and bacteria to multiply in parallel. The data also suggests the role of concomitant phage persistence mechanisms associated with delayed lysis of infected cells, such as carrier state infection. From an ecological and evolutionary standpoint this type of phage-host interaction is consistent with the Piggyback-the-Winner model, which suggests a preference towards lysogenic or other “benign” forms of phage infection when the host is stably present at high abundance.

**Teaser:** CrAss-like phage persistence in *Bacteroides* is associated with capsule phase-variation and additional unexplored mechanisms.

## Introduction

The crAss-like bacteriophages are a recently described family of dsDNA tailed bacterial viruses of the order Caudovirales that are predicted to infect bacteria of the phylum Bacteroidetes (*1*). They are present in variety of host-associated, aquatic and terrestrial habitats, but are especially predominant in the human gut (*2*–*4*). The presence of the prototypical crAssphage (p-crAssphage) in >70% of human gut metagenomes together with its exceptional abundance in some of the samples (>90% of the gut virome and >20% of total faecal DNA) led to its description as “the most abundant virus of the human body” (*4*). Furthermore, p-crAssphage was found to be widespread in the global human population with sequence variants correlating with geographic location (*5*). Based on the observation of related genomes in the non-human primate gut, it was proposed that p-crAssphage shares a long co-evolutionary history with the human species. In addition, limited dietary associations, but no link to disease was observed in crAssphage colonisation patterns (*3*). A recent longitudinal study of the human gut virome highlighted long-term (months and even years) high-level persistence of different combinations of crAss-like phages across individuals (*6*). When vertically transmitted from mothers to their infants or transplanted into in the course of faecal microbial transplantation (FMT), crAss-like phage engraft successfully and stably persist for many months (*7, 8*). CrAss-like phages can also persist in pure bacterial culture *in vitro* over many serial passages without any significant effect on the density of host population (*9*). The overall effect of stably persisting crAss-like phages, sometimes reaching absolute concentrations of 10^11^ genome copies per gram of faeces (*6*), on the structure (composition) and function of the gut microbiome is yet to be determined. Despite a lack of disease associations with p-crAssphage, correlations with other candidate genera and individual strains within the family of crAss-like phages has not been examined. Certain recent studies hinted at depletion of certain crAss-like viral clusters in an IBD patient cohort (*10*), while at the same time a genus VI crAss-like phage, termed IAS virus, was prevalent in HIV patients with unexplained diarrhoea (*11*).

It remains unclear what mechanisms are responsible for the long-term persistence and high prevalence of crAss-like phages in the human gut. Several mechanisms of phage persistence are possible, including spatial heterogeneity of microbial habitats in the gut (*12*), physiological stochasticity of phage sensitivity phenotype in their host bacteria (*13*), reversible genetic switching (phase variation) of surface receptor expression (*14*), constant sweeps of new mutations leading to arms race co-evolution with their hosts (*15*), or perhaps, an unusual life cycle (carrier state infection, pseudolysogeny) of the viruses themselves.

Establishing these mechanism(s) is a challenging task due to the lack of tools for genetic manipulation, and the low similarity of crAss-like phage genomes and proteins to well characterised model viruses (*16*). Our limited understanding of biological properties of crAss-like phages derives from crAss001, DAC15, and DAC17, the first three cultured human crAss-like phages (all members of the candidate genus VI) (*9, 17*).

Here, we report the first comprehensive effort to characterise interaction of crAss-like phages with their hosts, focusing primarily on crAss001. We demonstrate that phase variation of capsular polysaccharides creates dynamic sub-populations of sensitive and resistant *Bacteroides* cells, driving long term phage persistence. A yet unexplained mechanism is apparently responsible for a paused infection cycle and delayed release of phage progeny from a significant fraction of infected cells, a phenomenon that may represent a novel type of carrier state infection.

## Results

### CrAss-like phages are capable of long-term stable persistence in the mammalian gut and select for resistance in their bacterial host

We have previously demonstrated the long-term persistence of crAss-like phages in the human gut microbiome (*6*). The levels of crAss-like phage colonisation in ten subjects varied from <10% of the total faecal virome to nearly 100%. Eight of the ten subjects were stably colonised by one or several individual crAss-like phage strains.

Different crAss-like phage genera tended to colonise with distinctive relative abundances. High level colonisation with relative abundances approaching 100% of the virome were typical of genus I, whereas genus VI had much lower relative abundance, possibly reflecting differences in their host abundance, mode of replication, or the rate of emergence of resistance in the host strains (**Fig. 1a**). To reinforce this observation we followed the dynamics of crAss-like phage persistence in a monoxenic mouse model using a cultivable representative of the family, ΦcrAss001. Mice (n = 6) colonised with *B. intestinalis* alone served as a negative control. Within the first few days upon bacterial gavage, faecal *B. intestinalis* reached levels of 10^11^ cfu g^-1^ regardless of the presence of phage (p>0.05 in Kruskal-Wallis test with FDR correction for all time points except day 31). ΦcrAss001 propagated to much lower levels of 10^6^-10^8^ pfu g^-1^. However, this level also remained until the termination of the experiment at 136 days (**Fig. 1b**).

**Figure 1.**
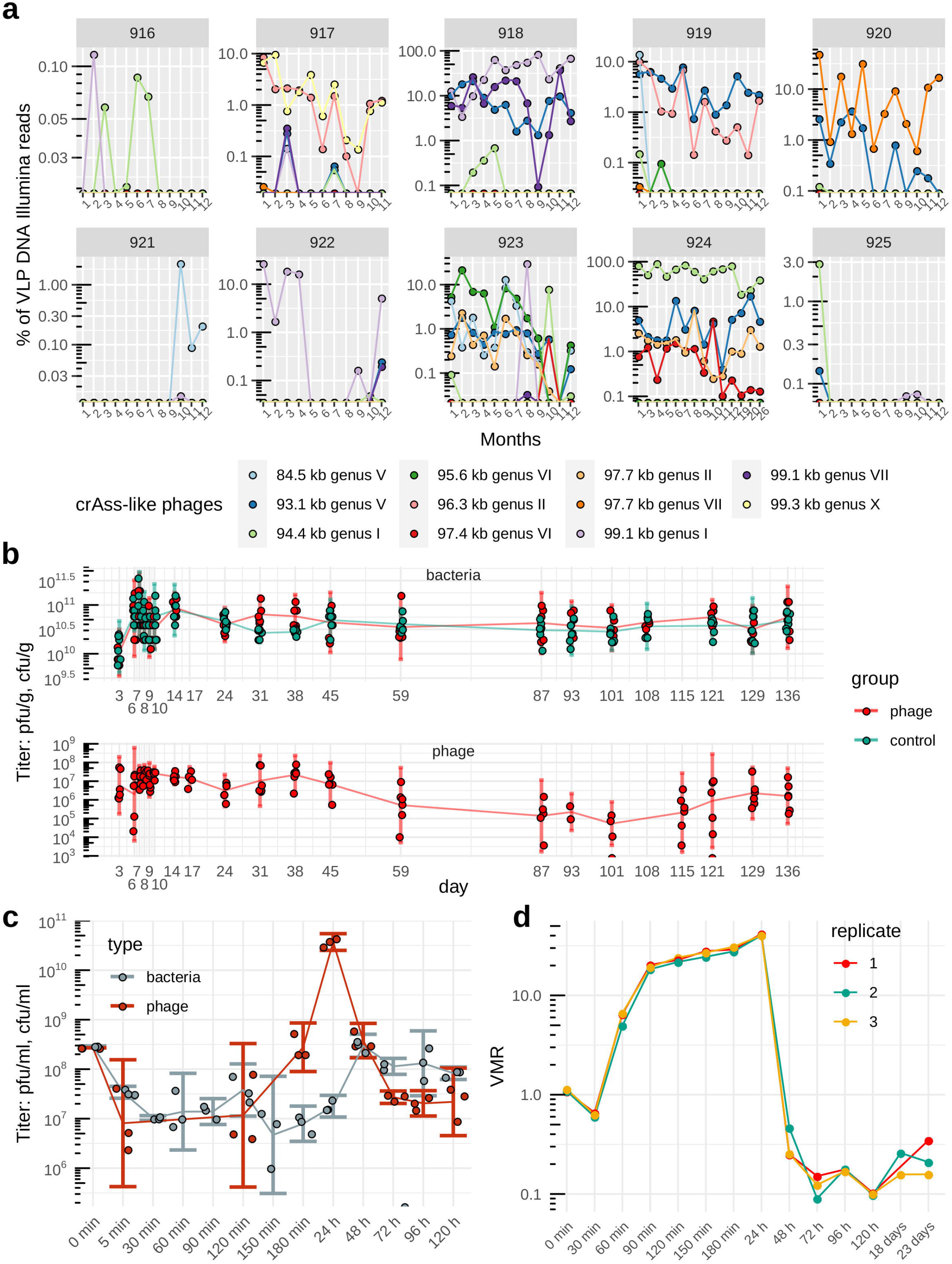
Long term persistence of crAss-like phages in the gut microbiome and *in vitro*. **(a)**, Persistence of 11 crAss-like phage genomes (belonging to candidate genera I,II,V,VI,VII,X according to Guerin et al. [2018] taxonomy) in faecal VLP fractions of 10 healthy adult individuals over a period of 12-26 months (*6*); **(b)**, Persistence of ΦcrAss001 in the gut of C57BL/6NTac mice (n = 6) mice colonised with ΦcrAss001/*B. intestinalis* APC919/174 mixture at an MOI=1; the control group of mice received oral gavage of *B. intestinalis* only; **(c)**, replication of ΦcrAss001 in exponentially growing culture of *B. intestinalis* (OD600=0.3 at infection with an MOI=1) followed by four daily transfers of phage/host co-culture in fresh broth; bacterial and free phage counts were obtained by plating and plaque assays respectively; **(d)**, VMR determined using shotgun metagenomic sequencing in the same phage/host co-cultures as in (c), with additional samples collected after 18 and 23 daily transfers.

Long term persistence of the phage-host pair quickly induces resistance to ΦcrAss001 in *B. intestinalis*. By day three from the first gavage, 11/17 (65%) of *B. intestinalis* faecal isolates were completely resistant to phage in spot and plaque assays, compared to 2% of spontaneously resistant cells in an overnight broth culture grown in the absence of phage (see below). By week twelve, 15/29 (52%) isolates were resistant. The sensitive isolates showed varied degrees of partial resistance that was manifested by formation of spots of variable turbidity. Unexpectedly, even bacterial isolates from the control animals showed a switch towards the phage-resistant phenotype in that while day three isolates were all sensitive (17/17), 8/29 (27%) isolates from week twelve were completely resistant, showing that a switch between sensitive and resistant phenotypes can occur in the host strain independently of phage attack.

This persistence can be recapitulated *in vitro*, in a *B. intestinalis* culture infected in its early logarithmic phase at multiplicity of infection (MOI) of 1, followed by overnight incubation and daily passages of resulting phage-bacterial co-culture. Initial absorption of phage resulted in 96.5% reduction of bacterial CFU after 30 min (**Fig. 1c**), followed by the first round of phage genome replication, which was largely completed by 90 min after infection resulting in a calculated virus-to-microbe ratio (VMR) of ∼22.8±1.2 (median±IQR, **Fig. 1d**). Parallel propagation of bacteria and phage for the first 24 hours maintained roughly the same level of VMR and resulted in high titre of phage particles 3.86×10^10^±7×10^9^ pfu ml^-1^ (**Fig. 1bc)**. Subsequent daily transfers led to a drop of VMR and phage titres, but resulted in increased viable bacterial titres and a phage/host dynamic that is stably maintained for as many as 23 daily transfers (**Fig. 1d**). Agreeing with previous observations in non-crAss-like phages and *Bacteroides thetaiotaomicron* (*14*), this evidence again points towards a rapid shift to dominance by resistant cells with retention of a sufficient sensitive sub-population to allow for low level phage reproduction and persistence.

### Resistance of B. intestinalis cells to ΦcrAss001 develops at high rate in vitro, but is reversible and associated with inhibition of phage adsorption

ΦcrAss001 forms turbid spots in soft agar overlays containing *B. intestinalis* APC919/174, confirming that bacterial growth can occur in the presence of phage (**Fig. 2a**). This could be a result of a high rate of lysogeny, or of a pre-existing resistant sub-population. resistant sub-population. To distinguish between these possibilities, transmission electron microscopy (TEM) was conducted on material collected from the center of the spot shown in **Fig. 2a**. Consistent with the hypothesis that a pre-existing resistant subpopulation of bacteria are present in the spot material, the electron micrographs revealed the presence of both uninfected and infected cells among the cell debris, including many instances in which cells in the process of division had phage progeny restricted to only one of the two nascent daughter cells (**Fig. 2b-f**).

**Figure 2.**
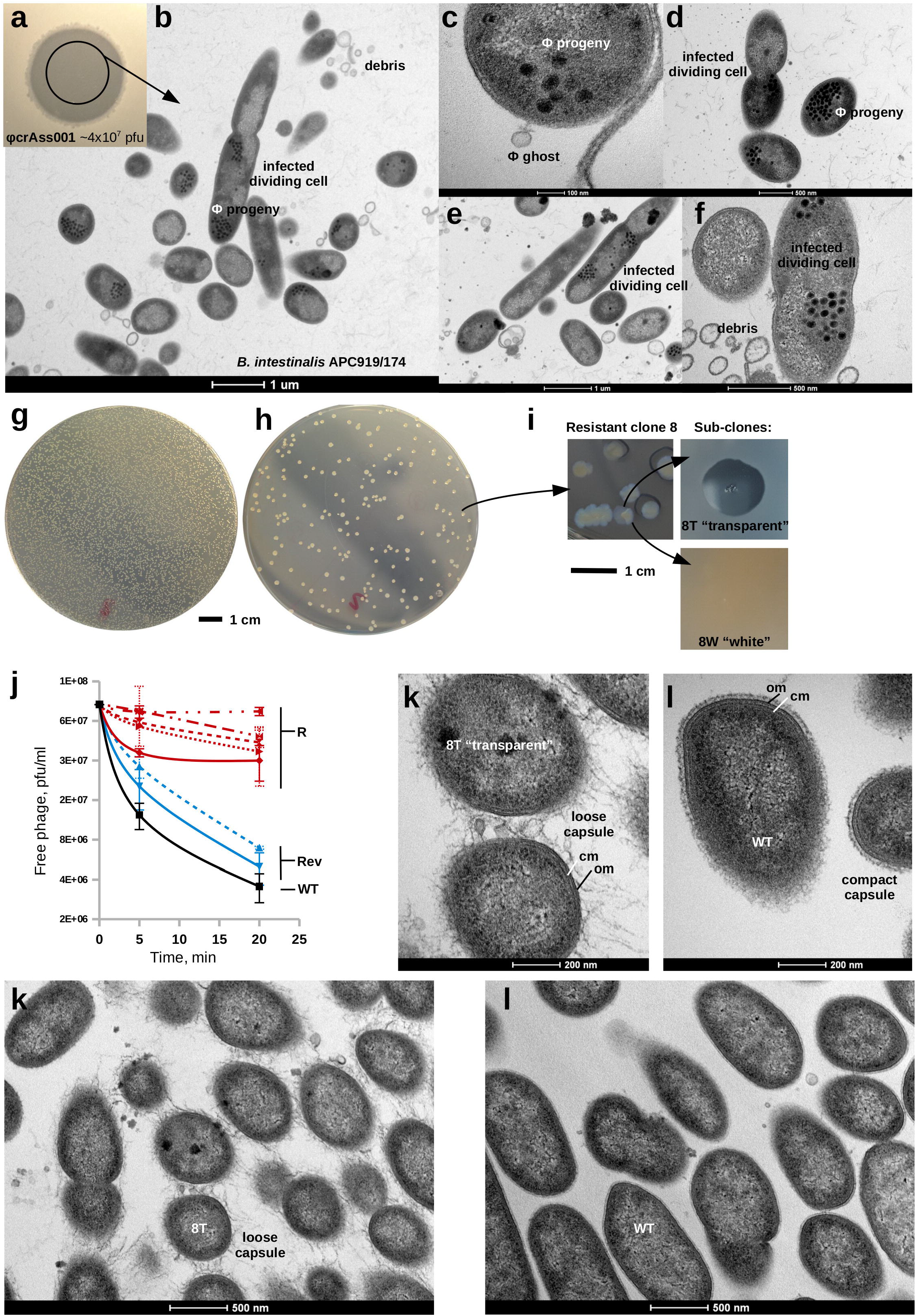
Resistance of *B. intestinalis* APC919/174 to ΦcrAss001 is reversible and associated with capsular polysaccharide alterations and loss of phage adsorption. **(a)**, spotting of 4×10^7^ pfu of ΦcrAss001 on *B. intestinalis* APC919/174 agar overlay results in turbid clearing zone; **(b-f)**, TEM (x9,900-105,000) of ultra-thin sections (80 nm) of samples taken from the centre of the spot shows the presence of un-infected cells amongst lysed cell debris, infected cells with empty virions (“ghosts”) still attached, and cells in the process of division with phage progeny visible inside of them; **(g) and (h)**, soft agar overlays inoculated with the same number of *B. intestinalis* CFU in the absence and presence of the excess of phage, respectively; **(i)**, phenotypic dissociation of colonies of phage-resistant clone 8, (8W, “white” – phage-sensitive subclone; 8T, “transparent” – phage-resistant subclone); **(j)**, kinetics of ΦcrAss001 adsorption to *B. intestinalis* phage resistant and sensitive clones; vertical axis, concentration of phage in the supernatant; horizontal axis, time passed after addition of phage; R, resistant clones (n=5); Rev, revertant sensitive sub-clones derived from one of the resistant clones; values are mean±SD from three independent experiments; **(k-l)**, ultrathin section TEM visualisation (x60,000) of altered cell surface morphology in phage-resistant clone 8T, compared to phage-sensitive WT strain; notable is the production of either loose or compact capsule; cm, cytoplasmic membrane; om, outer membrane.

In the presence of an excess of phage in soft agar overlays (efficiency of plating assay, EOP), clonal cultures of *B. intestinalis* APC919/174 had an immediate resistance rate of 2.06±0.14% (mean±SD, **Fig. 2g-h**), which agrees with 96.5% cells being infected above. None of the tested colonies contained DNA of ΦcrAss001 detectable by PCR. When ten resistant clones were each sub-cultured into 12 separate clones, four of the 120 (3.3%) reverted to sensitive phenotype. Resistant clones demonstrated a dramatically reduced ability to adsorb phage, while the spontaneously sensitive ‘revertants’ had restored phage adsorption (**Fig. 2j**). Some of the sub-clones had altered colony morphologies, with resistant colonies being more transparent, while sensitive ones were opaque (e.g. sub-clones 8T and 8W in **Fig. 2i**). When a resistant clone 8T was subjected to an EOP assay the rate of survival was raised to 88.7±3.3%, but in the sensitive sub-clone 8W it had returned closer to the original level of 11.3±0.9%. TEM of 8T cells revealed an altered “hairy” appearance, with a loose and bulky polymeric substance indicative of production of a different type of capsular polysaccharide (CPS) (**Fig. 2k**). In contrast, the wild-type (WT) culture demonstrates a much more compact thinner layer of capsular polysaccharide intimately attached to the outer membrane (**Fig. 2l**).

Together, these data suggests that cultures of *B. intestinalis* contain mixed populations of cells. The transient phage-resistant phenotype results from constantly ongoing reversible switching of expression of cell surface structures through some genetic or epigenetic mechanism. Approximately 2% of cells in naïve cultures were resistant to phage. Exposure to phage in continuous culture *in vitro* or in the murine gut drives the bacterial population towards phage resistance, but this resistance does not reach 100% or lead to the extinction of the phage under the conditions tested.

### Phase variation of multiple surface loci and concomitant capsular polysaccharide switching in B. intestinalis

In order to identify the mechanism of resistance we subjected ten resistant clones and two revertant sensitive sub-clones to whole genome sequencing. We also performed sequencing of samples collected from the long-term *in vitro* persistence experiment 5 min – 28 days following the inoculation and some of the faecal samples collected in long term murine colonisation experiment. For the wild type strain, resistant clone 8T (“transparent”) and its corresponding revertant clone 8W (“white”), complete circular genomes were assembled. A comparison of the three assemblies revealed that the only detectable differences consist of the inversion and rearrangement of several genomic loci (**Fig. S1a**). Similar to previous observations with B. *thetaiotaomicron* (*14*), *t*hree types of events were detectable, including, (a), large inversions involving entire gene clusters comprised of genes coding for TonB-dependent transporters (*18*) and/or RagB/SusD-like nutrient uptake proteins (*19*), often flanked with a dedicated integrase/recombinase gene; (b), a more complex type of reshuffling of clusters containing ABC-transporter genes (**Fig. S1b**); and (c), the inversion of small, <200 bp regions adjacent to capsular polysaccharide biosynthesis operons that contain potential promoters. Remarkably, all these genomic loci, designated as “phase variable regions” (PVRs), can be directly implicated in phage sensitivity. Their encoded products are either components of cell surface structures with potential to serve as receptors or barriers for phage adsorption or potential phage defence molecules (e.g. restriction/modification system).

Transcriptomic analysis of serial daily broth co-cultures of *B. intestinalis* and ΦcrAss001 1-5 days after the initial inoculation revealed an array of genes showing divergent responses to the presence of phage (p<0.01, DESeq2). The strongest transcriptional response in the host cells, already visible after the first 24 h, was associated with PVRs and consisted in the strong ∼100-fold repression of the PVR9 capsular polysaccharide (CPS) biosynthesis operon, the up-regulation of similar CPS operons associated with loci PVR7, PVR8 and PVR11 (**Fig. 3a**), and the up-regulation of a group of nutrient uptake genes in a complete inversion of the integrase/recombinase gene-flanked operon PVR3. Importantly, similar alterations of gene expression patterns relative to the phage-naïve WT culture of *B. intestinalis* can be observed in the resistant clone 8T (in the absence of phage). Its phage sensitive derivative 8W displays restored levels of PVR9 expression, down-regulated (compared to WT) PVR7, PVR8 and PVR11 and up-regulated expression of PVR3.

**Figure 3.**
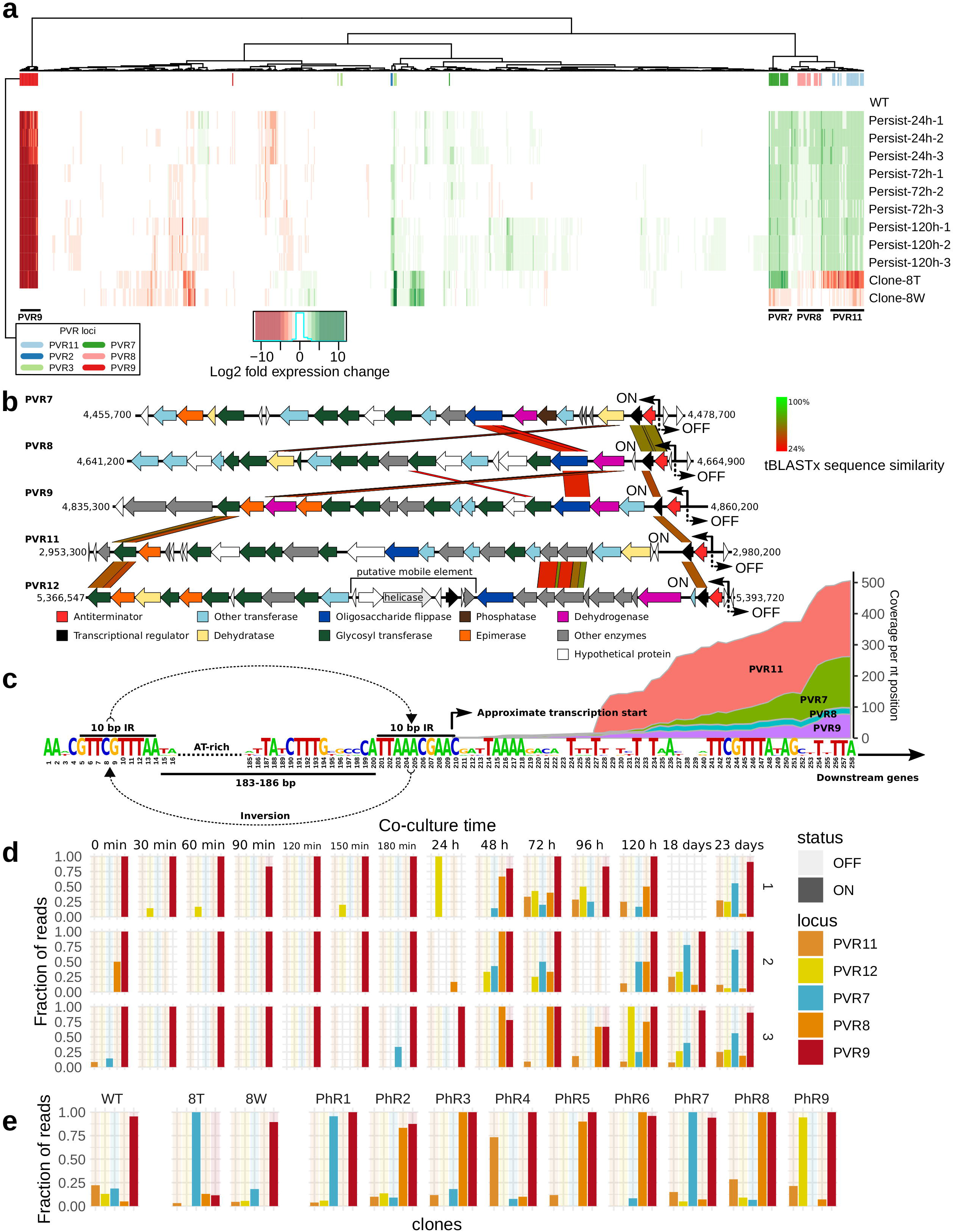
Structure and transcriptional control of five phase-variable capsular polysaccharide (CPS) operons (PVR7, PVR8, PVR9, PVR11, PVR12) in *B. intestinalis* APC919/174. **(a)**, host transcriptional response in a *B. intestinalis* APC919/174-ΦcrAss001 serial broth co-culture (24-120h *in vitro* persistence) experiment, analysed using RNAseq (values are log2 fold gene transcript expression change relative to the un-infected control from two independent experimental runs); clone-8T, phage resistant derivative of the parental strain; clone-8W, spontaneous phage-sensitive revertant clone; only genes with significantly changed expression (p < 0.01 in DESeq2, n = 826) are shown; genes, associated with phase-variable genomic regions (PVR) are marked with colour bars on top; **(b)**, structure of phase variable operons, encoding five different CPS; protein sequence homologies (tBLASTx) between gene products are shown as coloured parallelograms; two-sided arrows and ON/OFF labels mark positions of invertible promoters; **(c)**, consensus sequence of the regions flanking invertible promoters in CPS biosynthesis loci; approximate transcription start was identified though analysis of RNAseq data; stacked area charts show RNAseq read coverage of regions adjacent to the promoter in four CPS loci; **(d-e)**, fraction or Illumina concordantly-aligned read pairs supporting orientation of invertible promoters in either ON or OFF directions; WT, wild type strain; clones 8T, PhR1-9, phage-resistant derivatives; 8W, phage-sensitive revertant; P18d/P23d, samples in (e) were taken from the long term *in vitro* persistence experiment (also see Fig. S3 for similar data on the long term *in vivo* persistence).

The CPS-encoding loci PVR7-11 have broadly differing gene content with limited levels of amino acid homology between certain shared protein orthologues (e.g. oligosaccharide flippases, dehydrogenases, dehydratases and glycosyl transferases; **Fig. 3b**). All of these loci share a putative invertible, moderately conserved promoter sequence of 183-186 bp (**Fig. 3c**) flanked by a perfect, completely conserved 10 bp inverted repeat GTTCGTTTAA at which recombination most likely occurs. This sequence partially overlaps with inverted repeats ARACGTTCGTN flanking phase variable promoters controlled by a serine recombinase family, master DNA invertase (mpi) in *Bacteroides fragilis* (*20*). At least two homologs of this enzyme are encoded in the genome of *B. intestinalis* APC919/174 (NCBI accession QDO70032.1 and QDO69715.1). A search for similar patterns in the *B. intestinalis* APC919/174 genome revealed the presence of an additional, fifth, CPS locus (denoted PVR12) with identical inverted repeat flanking its promoter region. This locus, however, did not show significant up- or down-regulation of expression in our RNAseq assays. By mapping the RNAseq reads we located the approximate transcription start in PVR7-11 inside the proximal copy of the inverted repeat (**Fig. 3c**), confirming that the invertible regions are capable of acting as promoters.

To explore the rate and frequency of phase variation on the complete genome scale and to identify other potential PVRs, we performed local alignment of individual long DNA reads (obtained from WT culture, resistant clones 8T and PhR5, and corresponding revertant sub-clones 8W and PhR5-1) to the assembled circular genome reference, searching for cases where parts of the same long read align inconsistently, indicating either inversion or translocation. This analysis revealed a series of strong recombination hotspots present in every culture tested. The majority were associated with already known PVRs, with exception of a few novel finds, one of which is the previously mentioned fifth CPS locus PVR12 (**Fig. S2**). We then used mapping of short shotgun reads to allow the quantitation of inversions in the long term *in vitro* and *in vivo* persistence experiments, ten resistant clones and one revertant sub-clone. By aligning paired-end short reads we calculated the relative proportions of cells with two opposite orientations of the invertible promoters of PVR7-12 in different cultures of *B. intestinalis* APC919/174 (**Fig. 3d-e**). We found that the sensitive WT culture is characterised by PVR9 promoter mainly in the ON orientation, while PVR7, PVR8, PVR11 and PVR12 are mainly OFF. In agreement with the RNAseq data, the resistant clone 8T shows a switch of CPS expression from PVR9 towards PVR7, while the WT genotype is largely restored in sensitive revertant sub-clone 8W. Interestingly, other resistant clones showed the PVR9 promoter in mostly the ON orientation, combined with an ON state in one of the three other CPS-associated PVRs. Variable patterns of promoter activation with PVR9 being ON were detectable in the long term *in vitro* and *in vivo* persistence experiments (**Fig. 3d, Fig. S3**).

We suggest that in the *B. intestinalis*-ΦcrAss001 phage-host pair some of the CPS can be permissive to phage infection, and even serve as phage receptors (possibly PVR9), while some other can be protective (PVR7, PVR8, PVR11 and PVR12) or neutral. This is similar to what was observed for a large panel of non-crAss-like phages (*14*), as well as crAss-like-phages DAC15 and DAC17 in *B. thetaiotaomicron (17*). We conclude that phase variation of CPS expression leading to phage resistance occurs spontaneously, even in the absence of phage. As a result, a dynamic equilibrium between sensitive and resistant sub-populations is maintained, which can be shifted towards the resistance phenotype by phage-driven selection. However, a constant switching back to sensitivity provides the phage with a constant supply of sensitive cells, thus assuring its long-term persistence.

### Delayed release of ΦcrAss001 progeny leads to multiple echo phage bursts and potentially provides an infection escape mechanism

We previously reported a remarkably low apparent burst size of only 2.5 pfu per cell infected with ΦcrAss001 (*9*). This cannot be explained by failure of phage to infect the majority of cells as only ∼2% cells in naïve *B. intestinalis* cultures were phage-resistant due to phase variation. At the same time, >50 particles per cell are visible in TEM images from ultrathin (80 nm) cell sections (**Fig. 2b, d, f**) and at least 20 new copies of phage genome are produced in an infected culture per copy of bacterial genome after 90 min of infection (as shown by metagenomic sequencing, **Fig. 1d**), ruling out the possibility of extremely low progeny counts per infected cell. Further to that, the efficiency of centre of infection tests (EOCI), showed that 56.6±12.7% of cells infected at an MOI=1, and 59.16±27.9% infected at an MOI=10, were capable to give rise to infection centres (plaques) in *B. intestinalis* lawns. Electron microscopic observations of cultures infected at an MOI=1 reveal that >90% of cells show signs of early stages of virion assembly process 40 and 90 minutes after infection (**Fig. S4**). Therefore, the observed low burst size cannot be explained by phase variation-based resistance or low progeny per cell counts and must from an unknown mechanism limiting replication and/or release of phage progeny from a significant fraction of infected cells.

In order to get insights into this mechanism, we performed strand-specific RNAseq analysis of a single phage replication cycle in *B. intestinalis* cells infected at an MOI of 1. Three transcriptional modules can be observed in the ΦcrAss001 genome, which also correlate with ORF orientation and the putative operon organisation of the genome (**Fig. 4a**). The expression of the early genes, located on the right end of the linear phage genome, is initiated between 0 and 10 minutes after phage infection. The predicted products of these genes include a transcriptional regulator, two conserved domain proteins with unknown function and a TROVE domain protein (possible ribonucleoprotein component). A recent study in a distantly related marine crAss-like phage Φ14:2 infecting *Cellulophaga baltica* indicated that a similarly organised operon of early genes is transcribed by a giant virion-associated multi-subunit RNA polymerase [RNAP, (*21*)], ejected into the cell in the course of infection (*22*). A homologue of this giant polymerase is present in all crAss-like phages and is also encoded by three large ORFs located in the central part of ΦcrAss001 genome (*4, 9, 23*). Proteomic analysis of the crAss001 virion also indicated the presence of the subunits of the giant polymerase in the assembled phage particle (*9*).

**Figure 4.**
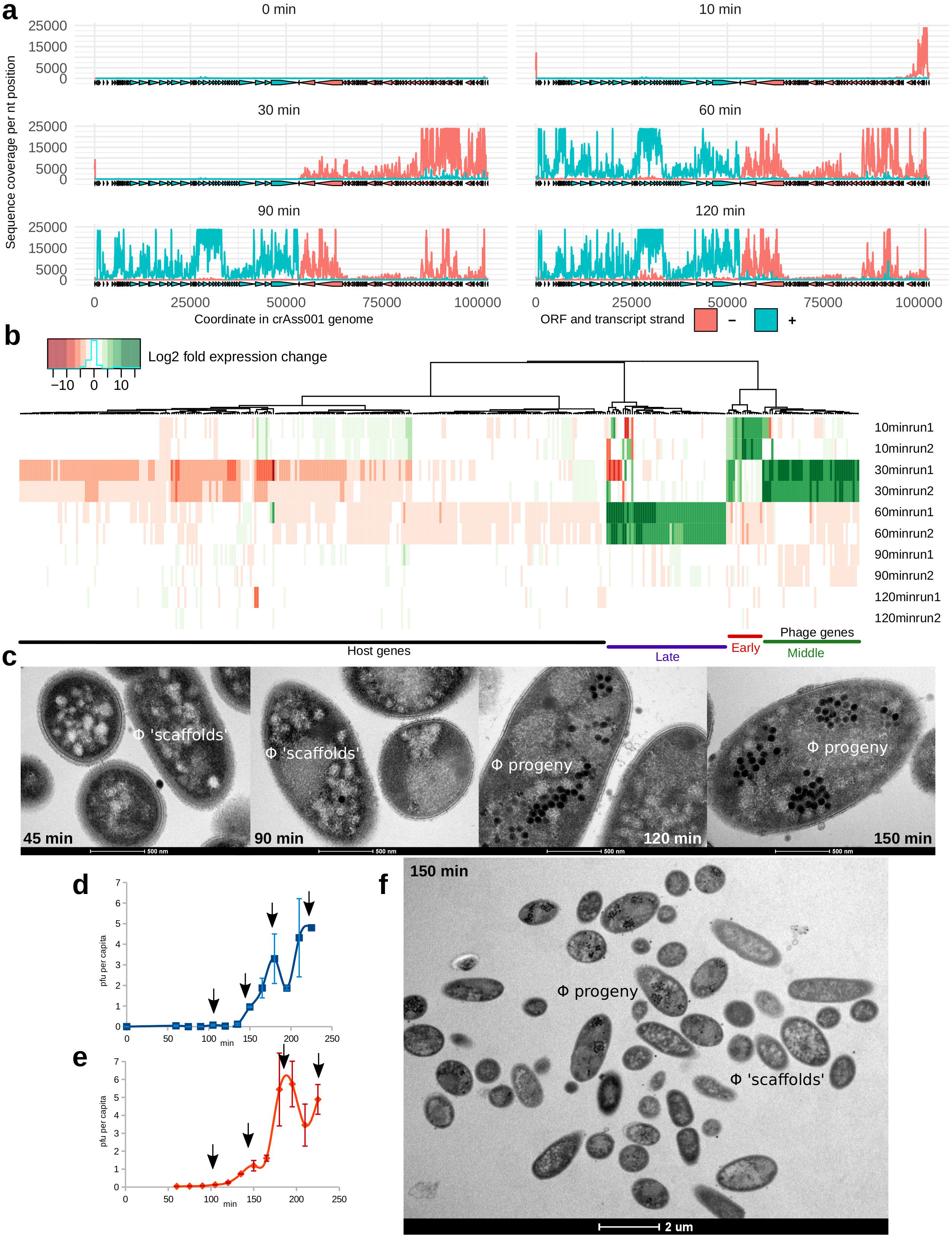
Transcriptional and morphological presentation of ΦcrAss001 infection in a one-step growth experiment. **(a)**, transcription of phage genome 0-120 min after infection revealed using stranded RNAseq; positive and negative strand ORFs and RNAseq coverage levels are shown in blue and red, respectively; plots are representative of two independent experiments; **(b)**, time course of transcriptional responses of host and phage genomes (values are log2 fold gene transcript expression change relative to the previous time point from two independent experimental runs); only genes with significantly different expression of sense DNA strand between any of the successive time points (p < 0.05 in DESeq2, n = 372) are shown; **(c)**, time-course of morphological changes in *B. intestinalis* cells infected with ΦcrAss001, ultrathin section TEM (x43,000); notable are intracellular phage progeny and possible “phage scaffolds” in the early stages of virion assembly; **(d-e)**, enumeration of per capita phage progeny produced in a one-step growth experiment by *B. intestinalis* cells infected with ΦcrAss001 [MOI=1 in panel (d), MOI=10 in panel (e)]; **(f)**, intact cells after first phage burst (150 min), still showing hallmark features of the early and middle infection stages (x6,000).

Transcription of the middle and late genes is initiated between 10-30 and 30-60 minutes, respectively, and is likely to depend on the action of the host RNAP [as suggested for Φ14:2 in (*21*)]. The middle genes are largely composed of functions associated with DNA replication and recombination, as well as nucleotide metabolism. Other genes of the same class can be involved in the degradation of host DNA and mRNA (HKD family nuclease, HicA mRNAse). The late genes, occupying the left-hand half of the genome, mainly encode structural proteins of the virion head and tail, as well as proteins participating in the virion assembly (terminase, chaperone) and cell lysis (holin, endolysin). Large ORFs coding for virion-associated RNAP subunits are located in the approximate centre of the linear genome and are transcribed during middle (gp49, predicted catalytic subunit, and gp50) or late (gp47) transcriptional stages. It is unclear why the temporal separation in RNAP subunit production might be needed. It may be that gp49 and gp50 are involved in transcription of the late genes in the same infection cycle in which they are produced, whereas gp47 is needed solely for the early genes transcription in the next cycle, or plays a role in the packaging, ejection or activation of the RNAP holoenzyme.

Interestingly, there was significant overlap in transcription of early, middle and late genes: once activated, transcripts of each of the three modules persisted through the rest of the replication cycle (**Fig. 4a-b**). The host transcriptional response was largely confined to strong genome-wide inhibition of transcription, occurring between 10-30 minutes preceded by the activation of certain energy metabolism functions at 10 minutes (most notably citrate synthase and NADP−dependent iso-citrate dehydrogenase, **Fig. 4b**).

Morphological changes in the infected cells were visible as early as after 45 minutes as the formation of variable-sized electron-light spots (**Fig. 4c**). Ninety minutes into the infection cycle, well after the onset of expression of structural genes, the first intact phage particles (electron-dense spots) begin to appear amongst the electron-light spots. Large clusters of viral particles are visible inside many of the infected cells at 120 minutes. Agreeing with the one step growth curves (**Fig. 4d-e**), cell lysis becomes noticeable at 150 minutes and widespread at 180 minutes. Interestingly, however, a large fraction of cells at 150 minutes remain intact, and only show signs of the early stage of infection with no properly formed phages particles (**Fig. 4f**). This observation can be interpreted as de-synchronisation of the phage replication cycle with some of the cells falling behind the others in terms of infection stage. Supporting this hypothesis is the continued expression of early and middle operon genes, overlapping in timing with late genes. This is further reflected in the one step growth curves, where at an MOI of 1 or 10, a series of discrete, evenly spaced bursts are apparent with a low total progeny output (∼3-6 pfu per capita).

Recently, bacteriophages similar to ΦcrAss001 in morphology and genome organisation (**Fig. S5a-b**) were isolated against *B. thetaiotaomicron* (*17*). Phages DAC15 and DAC17 are also capable of long-term persistence in serial broth cultures of *B. thetaiotaomicron* VPI-5482 (**Fig. S5c**) and do not significantly reduce the bacterial population. While the wild-type strain produce eight different phase-variable CPS, a panel of engineered mutants have been developed, which included an acapsular version, as well as eight strains expressing only one of the eight different CPSs, referred to hereafter as CPS1-8 (*24, 25*). Phages DAC15 and DAC17 were reported to infect the *cps3*^+^ mutant efficiently, while only showing weak replication in WT VPI-5482 and other single-CPS expressing derivatives, indicating that CPS3 plays a permissive role in phage infection.

In our hands, both DAC15 and DAC17 were able to persist stably in serial daily broth cultures of *cps3*^+^, as well as in WT (albeit at lower levels, **Fig. S5d**). This is despite the fact that an inability to switch to expression of alternative CPSs leads to a very low fraction of pre-existing resistant cells (< 10^−5^) in populations of *cps3*^+^, compared to 52.8±1.2 – 64.2±3.0% of cells resistant to DAC15/ DAC17 in the WT strain. This indicates that the removal of CPS variability leads to greatly reduced resistance to phage but does not eliminate the ability of phage to persist long-term. In one step growth experiments at an MOI=1, DAC15 and DAC17 behaved very similarly to ΦcrAss001, producing a series of bursts and a low apparent output of progeny (**Fig. S5e-f**). However, unlike in *B. intestinalis-*ΦcrAss001 system, infection of *cps3*^+^ mutant at MOI=10 resulted in a single burst of ∼160 pfu per capita, a value which is in a much closer agreement with the progeny size observed microscopically (**Fig. S5g**).

These data suggests that a mechanism of gradual release of phage progeny operates in *Bacteroides* cells infected with crAss-like phages ΦcrAss001, DAC15 and DAC17. This mechanism is largely independent of the dynamic phase variation mediated resistance discussed above. It appears that at high MOI the mechanism might become overwhelmed, or that the stress response in cells simultaneously infected by multiple phage particles switches it off. This leads to abrupt cell lysis and release of large phage progeny simultaneously, in a “classical” virulent phage manner.

## Discussion

Since their discovery in 2014, crAss-like phages have become one of the most intriguing elements of the human virome. Based largely on metagenomic observations, this group of tailed dsDNA bacteriophages differs significantly from other members of the order Caudovirales in terms of their genome organisation, gene content, virion morphology, life cycle and ecological distribution (*3, 6, 9, 16*). Of special interest is their unexplained, overwhelming prevalence in the gut microbial community, their ability for long term persistence and seemingly benign interaction with their hosts (*6*). The current, provisional classification of crAss-like phages based on gene sharing patterns, lists ten genera (denoted I-X) in four subfamilies (*Alphacrassvirinae, Betacrassvirinae, Gammacrassvirina*e and *Deltacrassvirinae*). Genus I includes the original p-crAssphage and is by far the most commonly detected and the most abundant member of the family in the human gut, at least in the Western populations (*23*). However, the only genus that contains a cultured representative is candidate genus VI (*Betacrassvirinae*) which are widespread but only moderately abundant in the human gut. Members of this genus were isolated from human faeces (ΦcrAss001), wastewater effluent (DAC15 and DAC17), and sea water (Φ14:2), infecting *B. intestinalis* (*9*), *B. thetaiotaomicron* (*17*) and *C. baltica*, respectively (*22*). All these phages were able to lyse target cultures at least partially and produce spots and plaques in a semi-solid agar overlay assay. Phage of genus I and other genera common in the human gut could be grown *in vivo* after transfer into oligoxenic mice or after faecal microbial transplantation of crAssphage-rich microbiota into human patients, and *ex vivo* as part of a native faecal community in a bioreactor, but have not as yet been isolated in pure bacterial cultures (*6, 7, 16, 26*).

We hypothesize that crAss-like phage-host systems encompass a range of behaviours, from one resembling virulent phages that produce plaques but whose replication in pure host cultures and complex microbial communities is typically supported at lower levels; to one similar to temperate phage that produce no plaques but persist in host populations at very high levels. It is notable that, regardless of the degree of virulence of a crAss-like phage, their stable and high-level persistence *in vitro* and *in vivo* has no obvious detrimental effect on the population densities or fitness of their bacterial host. This phenomenon is generally consistent with a temperate mode of replication, but we conclude from our analysis of ΦcrAss001, DAC15 and DAC17 that these viruses are non-lysogenic. Specifically, (i) no cases of related prophages have ever been observed in public databases of bacterial genomes; (ii) no clear lysogeny modules could be identified their genomes; and (iii) lysogens or pseudolysogens could not be obtained experimentally.

Observations of long-term persistence of virulent bacteriophages in the mammalian gut are not unprecedented. Experiments in monoxenic mice revealed that administration of virulent phage fails to lower the levels of target *E. coli* bacteria and often leads to continuous shedding of high phage levels in faeces (*27, 28*). Alternative mechanisms of non-lethal, persistent forms of infection were described in virulent phages, including the carrier state and unstable pseudolysogeny in *Campylobacter jejuni, Salmonella typhimurium* (phage P22) and *Pseudomonas aeruginosa* (*29*–*31*). In *C. jejuni*, virulent bacteriophages CP8 and CP30A have been shown to maintain equilibrium with their hosts for many generations and to persist intracellularly in an episomal form without integration into the host chromosome. It has also been demonstrated that the carrier state can have a profound effect on bacterial host physiology, such as a major acquisition of self-targeting CRISPR spacers, decreased fitness, loss of colonisation potential and motility in *C. jejuni* (*32*). More recently a carrier state was described in *P. aeruginosa* infected with ssRNA levivirus LeviOr01 (*33*).

Infection with LeviOr01 caused dissociation of the sensitive host into two sub-populations: a small subset of phage-carrying superinfection immune cells, spontaneously and continuously releasing phage progeny, and a larger subset of uninfected non-carrier cells. Together, previous observations suggest that unconventional forms of phage infection span a wide taxonomic variety of bacteria and levels of complexity of phages (from *Leviviridae* containing only four genes in their genomes, to *Myoviridae* with hundreds of genes).

The genus *Bacteroides* comprises strictly anaerobic Gram-negative non-sporeforming bacteria, many of which are predominant members of the human gut microbiome and are highly adapted to the gut environment (*34*). Genome plasticity, extensive regulation of gene expression and high adaptability to changes in their environment are characteristic features of this bacterial group (*35*). Phase variation of surface structures resulting from DNA inversions catalysed by serine and tyrosine recombinases has been implicated as an important mechanism in *Bacteroides* for adaptation to environmental change, improved fitness, and efficient colonisation of the mammalian gut (*20, 24, 36*–*38*). Recently, phase variation of multiple surface structures including CPS, S-layer lipoproteins, TonB-dependent nutrient receptors and OmpA-like proteins, has also been implicated in dynamic switching of diverse phage resistance/sensitivity patterns in *B. thetaiotaomicron* (*14*). Furthermore, crAss-like phages DAC15 and DAC17 were shown to preferentially infect a phase variant of *B. thetaiotaomicron* expressing a specific type of CPS (*17*). Recent modelling with phase variable phage receptor in *Haemophilus influenzae* demonstrated the role of phase variation in bacterial “herd immunity”, a phenomenon in phage-host population dynamics where “protective quarantining” of one third of the host cells already confers significant protection against phage attack to the whole population (*39*).

In this study we shed some light on the mechanisms underpinning persistence of crAss-like phages using the ΦcrAss001-*B. intestinalis* pair as a model. We observed that, similarly to metagenomic findings from the human gut (*6, 8*), a monoxenic mouse model was able to support long term and stable persistence of ΦcrAss001. Bacterial clones recovered from colonised mice showed varied degree of resistance to phage, but no obvious cost with regards to bacterial population density was associated with phage colonisation. We propose that that at least two separate mechanisms operating simultaneously are responsible for observed persistence *in vitro*: (i) dynamic and reversible acquisition of phage resistance in the host population, dependent on phase variation of CPS; (ii) delayed release or progeny from infected cells, resulting in a pseudolysogenic-like or carrier state phenotype. (**Fig. 5**).

**Figure 5.**
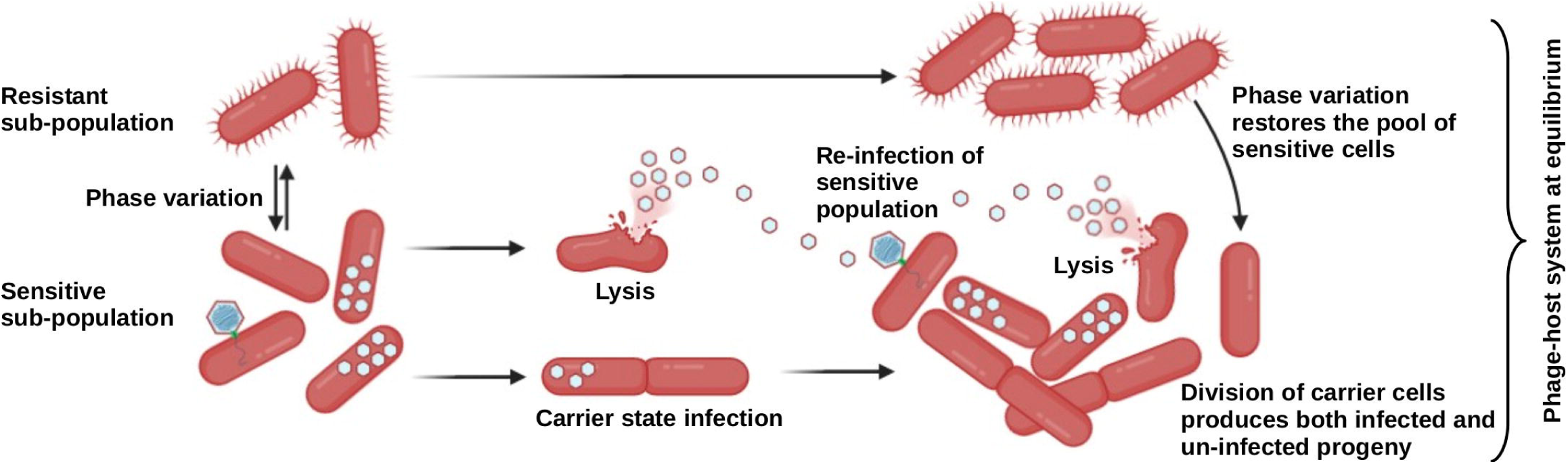
Model of perpetuated replication of crAss-like phage in *Bacteroides* cultures. (Created using BioRender.com). See Discussion for details.

Sustained phage attack resulted in either switching off the PVR9 CPS locus, implicating this structure in phage adsorption, or increased expression of alternative CPS’s (PVR7, PVR8, PVR11 and PVR12) with possible protective effects. We propose that the dynamic switching between different CPS types maintains an equilibrium of sensitive and resistant cells, allowing for restricted (“quarantined”) phage replication and enabling long term phage persistence and herd immunity in the bacterial population. While phage attack provides an obvious selective force driving switching to resistance states, the force driving a return to sensitivity remains unidentified. In complex microbiota conditions, expression of CPS protective against one phage could make cells sensitive to another phage present in the same environment (*14*), creating an equilibrium of CPS expression states driven by opposing phages. In monoculture, however, such diverse phage selective forces are absent, making the metabolic cost of expression of a particular protective CPS an obvious reason for switching it off. Availability of a single-CPS mutant panel in *B. thetaiotaomicron* prompted us to check whether persistence of DAC15 and DAC17 is still possible despite inability of the strain to evade phage by swapping its CPS types. Indeed, not only persistence in such strain was still possible, but actually occurred at higher levels than in the WT.

We hypothesise that a second mechanism of phage persistence operates in parallel with herd immunity. This is based on the low apparent burst size (despite progeny counts of >50 per cell under the EM), multiple sequential smaller bursts and cells carrying phage progeny without lysing in a timely manner, the high expression of mainly early genes (regulatory proteins) and some of the middle genes (replication functions) in broth cultures of *B. intestinalis* in the presence of phage. Several explanations are possible: (a) carrier state, persistence in the form of fully formed or incompletely formed viral particles in the host cytoplasm; (b) unstable pseudolysogeny, persistence of phage genomes in an episomal form; (c), inhibition of phage reproduction by cellular mechanisms and curing of infected cells. Interestingly, overwhelming the cells with high MOI was enough to overcome this mechanism of gradual release of progeny in *B. thetaiotaomicron* but not in *B. intestinalis*.

We suggest that the interplay between phase variation of phage receptors and the carrier state infection enable highly synchronised replication of a crAss-like phage with its host at a population level, and leads to an equilibrium of phage/host ratio maintained over multiple generations. It has been proposed that high availability of bacterial prey in aquatic environments and at mucosal surfaces selects for temperate bacteriophage lifestyles and lysogenic mode of replication. Such persistent infection is highly beneficial for the phage and is at least ‘affordable’ (if not also beneficial) to the host (*40, 41*). This ecological model has become known as “piggyback-the-winner”, as opposed to the “kill-the-winner” model (*41*). The mammalian gut presents an example of one of the most densely populated microbial communities, with *Bacteroides* being one of its most abundant and temporally stable bacterial genera (*6, 42, 43*). Our expectation was that the majority of bacteriophages in such an environment would be temperate, “piggybacking” on the ecological success of their bacterial hosts (*44*–*46*). At the same time our recent metagenomic analysis highlighted the prevalence of apparently virulent bacteriophages in the healthy core virome (*6, 10*). One could hypothesise that crAss-like phages, incapable of classical lysogeny and comprising a large part of these metagenomic sequences, employ alternative life cycle strategies, such as the one highlighted in this this study in order to benefit from “piggyback-the-winner” dynamics.

While the ecological benefits of carrier state infection are obvious for the phage, we can expect that the bacterial host party should also derive certain benefits from this interaction. In addition to exerting a constant selective pressure leading to host diversification through phase variation and point mutations and an improved overall fitness, persistent phage can participate in lateral gene transfer, provide protection from incoming competitor strains and superinfection immunity against cognate phages.

## Acknowledgements

This research was conducted with the financial support of Science Foundation Ireland (SFI) under Grant Number SFI/12/RC/2273, a Science Foundation Ireland’s Spokes Programme which is co-funded under the European Regional Development Fund under Grant Number SFI/14/SP APC/B3032, and a research grant from Janssen Biotech, Inc. Authors wish to thank Frances O’Brien and Tara O’Driscoll (UCC) for their technical help with germ-free mouse experiment.

## Author contributions

Conceptualization, A.N.S. and C.H.; Methodology, A.N.S., E.V.K., N.S., A.J.H., C.Heuston and D.S.; Software, A.N.S.; Formal analysis, A.N.S., E.V.K., D.S. and C.Hill; Investigation, A.N.S., E.V.K., C.Heuston and S.S.; Resources, A.J.H., D.S. and C.Hill; Writing – original draft, A.N.S. and E.V.K.; Writing – review and editing, A.N.S., E.V.K., A.J.H., C.Heuston, D.S., R.P.R. and C.Hill; Supervision, D.S., C.Hill and R.P.R.; Project administration, C.Hill; Funding acquisition, C.Hill and R.P.R.

## Data and code availability

All data needed to evaluate the conclusions in the paper are present in the paper and/or the Supplementary Materials. Further information and requests for data and resources should be directed to and will be fulfilled by Dr. Andrey Shkoporov. Raw sequencing data are available from NCBI databases under BioProject PRJNA678472. Raw numerical data and images used to produce **Fig. 1-4** are available from the Supplementary Dataset (https://figshare.com/s/c3485a7633d3632579ca).

## Materials availability

Phage ΦcrAss001 and its host *B. intestinalis* APC919/174 are available from DSMZ culture collection under catalogue numbers DSM 109066 and DSM 108646 respectively. Phages DAC15 and DAC17, as well as *B. thetaiotaomicron* VPI-5482 capsule mutants are available on request from Prof. A.J. Hryckowian.

## Materials and methods

### Bacteriophage ΦcrAss001 and related viruses

The host strain *B. intestinalis* APC919/174 was propagated anaerobically on Fastidious Anaerobe Broth (FAB, Neogen, UK) and agar (FAA) at 37°C as described before (*9*). To generate high titer phage lysates, early log phase culture (OD_600_=0.2; ∼2×10^8^ cfu ml ^-1^) were infected with ΦcrAss001 at an MOI of 1. Cultures were left to grow overnight at 37°C in anaerobic jars, centrifuged for 15 min at 5,200 g and 4°C and filtered through 0.45 μm polyethersulfone (PES) syringe-mounted filter membranes. Plaque and spot assays throughout different experiments were performed after filtering 1 mL of phage/bacteria co-cultures, performing serial dilutions and combining 100 μl of them with 300 μl of a fresh APC919/174 overnight culture in 0.4% bacto agar overlays. Incubations were performed anaerobically at 37°C. Bacteriophages DAC15 and DAC17 and their host strains *Bacteroides thetaiotaomicron* VPI-5482 (WT and *cps3*^*+*^) were propagated in the same conditions as *B. intestinalis*/ΦcrAss001.

### Germ-free mouse model

The germ-free animal experiment was performed in accordance with the European Communities Council Directive 2010/63/EU under an authorisation issued by the Health Products Regulatory Authority (HPRA, Ireland), with approval from the Animal Experimentation Ethics Committee (AEEC) of University College Cork. Twelve male 13 week old (26-29 g) germ-free C57BL/6N mice (RRID:MGI:5651595, original breeding stock from Taconic, USA; bred in house) were housed in groups of 2-3 mice in HEPA filtered individually ventilated cages to maintain germ-free status. All dosing and faecal sampling occurred under sterile conditions in a biosafety cabinet to ensure there was no confounding colonisation. Group 1 (n = 6) were orally dosed on three successive days with 100 μl of phage/bacteria suspension (10^9^ cfu of bacteria, 2×10^8^ pfu of phage per gavage). Group 2 were orally dosed with bacteria only (10^9^ cfu per gavage). Faecal samples were collected for 146 days after the first dose (days 3, 6-10, 14, 17, 24, 31, 38, 45, 59, 87, 93, 101, 108, 115, 121, 129, 136, 146) and subjected to plating and plaque assays for enumeration of bacteria and phage (**Fig. 1b**). On the last day of the experiment animals were humanely euthanised by cervical dislocation.

### In vitro experiments with crAss-like phages

Phage adsorption experiments (**Fig. 2j**) were conducted in aerobic conditions at room temperature on cells from an overnight *Bacteroides* culture (10 mL, optical density adjusted to OD_600_=0.2), triple washed with growth medium and resuspended in the same media volume before the experiment. Phage was applied at an MOI=1, aliquots were removed, filtered as described above and subjected to plaque assays 5 and 20 min after addition of phage.

Efficiency of bacterial plating (EOP) in the presence of phage (**Fig. 2g-h**) was conducted as follows: Serial tenfold dilutions of overnight *Bacteroides* cultures were inoculated into 3 mL of semi-solid FAA (0.4% agar) with or without addition of 100 μL high-titre phage (∼10^10^ pfu mL^-1^) and poured onto 100 mm diameter FAA agar plates. Efficiency of lysogeny was determined as a percentage of colonies observed on phage-containing overlays relative to the total counts on negative control overlays after 48 hours of anaerobic incubation at 37°C.

*In vitro* phage persistence experiment (**Figs. 1c-d, 3a, 3d, S5d**) was performed as follows: ten millilitre liquid cultures (n=3) of *Bacteroides* were infected at OD_600_=0.2 and MOI=1, and incubated overnight. The co-culture was then transferred daily, for five days in fresh FAB medium at 1:50 ratio.

One-step growth curves (**Fig. 4d-e, S5e-g**) were built as follows: early logarithmic phase (OD_600_=0.2) cultures of *Bacteroides* (20 mL) were infected at an MOI of 1 or 10 for 5 min at room temperature, followed by centrifugation at 5,200 g, 4°C for 15 min, removal of supernatant and re-suspending of the infected cells in fresh FAB medium. Incubation was continued anaerobically at 37°C for further 225 min with removal of 1 mL samples every 15 min. Samples were filtered through 0.45 µm pore PES filters and subjected to standard plaque assays with appropriate dilutions. Efficiency of centre of infection (EOCI) was determined by doing plaque assays with infected cells at the same time intervals but without filtration step.

### Transmission electron miscroscopy

Ultrathin section TEM of *Bacteroides* cells grown in broth cultures (**Fig. 2k-l**) or embedded in semi-solid agar (**Figs. 2b-f, 4c, 4f, S4**) were performed as follows. Pellets from 1 mL broth cultures or agar overlay fragments (∼100μL volume) were immediately fixed and stored in 1 mL of 2% (v/v) glutaraldehyde, 1.5% (w/v) parafolmaldehyde, 75 mM Tris-HCl pH 7.5 solution. Samples were post fixed with 1% Osmium tetroxide in 0.1 M Sorenson’s phosphate buffer for 1 hour at room temperature. A series of graded dehydrations were performed with increasing ethanol concentrations (30%, 50%, 70%, 100%; cells from broth cultures were pelleted by centrifugation between steps).

Dehydrated samples were embedded in Epon resin. Briefly, dehydrated samples were transferred into 100% acetone and then 50:50 acetone:Epon resin for a minimum of 1 hour. Samples were then put in 100% Epon resin at 37°C for 2 hours. Resin was polymerised at 60°C for a minimum of 24 hours. Embedded samples were sectioned at 80 nm thickness onto copper grids using a Leica UC6 ultramicrotome. Samples were stained with 2% uranyl acetate for 20 minutes and 3 % lead citrate for 5 minutes. Grids were then imaged on a FEI Tecnai 120 at 120 kV accelerating voltage. TEM of purified phage particles (**Figs. S5a-b**) was performed as described before (*9*).

### Shotgun sequencing and analysis of bacterial genomes

Shotgun sequencing on Illumina platform was performed for parental strain APC919/174 (**Figs. 3b, 3e, S1, S2**), resistant clones PhR1-9, 8T and spontaneous sensitive revertant derivative (8W) of the latter (**Figs. 3e, S1, S2**), as well as samples from *in vitro* phage persistence experiment and faecal samples from mice colonized with ΦcrAss001/*B. intestinalis* phage-host pair (**Figs. 3d, S3**). BioSample accession number are SAMN16802447-SAMN16802559 and SAMN16803214. Genomic DNA was extracted from bacterial cultures and phage-host co-cultures using DNeasy Blood & Tissue Kit (Qiagen) in accordance with the manufacturer’s Gram-negative bacteria protocol. Faecal DNA was extracted using QIAamp Fast DNA Stool Mini Kit (Qiagen). Concentration of DNA was determined using the Qubit dsDNA HS kit and the Qubit 3 fluorometer (Invitrogen/ThermoFisher Scientific). One hundred nanograms of purified DNA sample were sheared with M220 Focused-Ultrasonicator (Covaris) applying the 350 bp DNA fragment length settings (peak power 50 W, duty factor 20 %, 200 cycles per burst, total duration of 65 s). Random shotgun libraries were prepared from genomic DNA using TruSeq Nano Library Preparation Kit (Illumina, cat #20015964) with dual-indexing following the standard manufacturer’s protocol. Paired-end sequencing with 2×150 nt chemistry was performed on HiSeq 4000 platform at Eurofins Genomics.

The quality of raw sequences (SRA database records SRR13062103-SRR13062215) was assessed using FastQC v0.11.5. TruSeq adapters were removed with cutadapt v2.4. To trim sequences and remove low quality reads, Trimmomatic v0.36 was applied using the following parameters: ‘SLIDINGWINDOW:4:20 MINLEN:60 HEADCROP:10’, yielding 2.2-9.7 million reads per sample (median 5.9 million).

For long read Oxford Nanopore sequencing genomic DNA from APC919/174, 8T, 8W, Phr5 and Phr5-1 (sensitive derivative of the resistant clone Phr5) was extracted as follows. Cells from two mL of overnight cultures were collected by centrifugation at 5,000 g for 10 min, washed by resuspending in 1 mL of TES (50 mM NaCl, 100 mM Tris-HCl, 70 mM disodium EDTA, pH 8.0) and re-centrifuged. After resuspending cell pellets in 0.4 mL of TES supplemented with 25% w/v sucrose and 30 mg mL^-1^ of lysozyme, 50 U of RNase I (Fisher Scientific) were added and samples were incubated at 42°C for 1 hour. Cells were then lysed by addition of 20 μL of 10% (w/v) sarkosyl and 5 μL of 20 mg mL^-1^ proteinase K solutions with incubation at 37°C until complete clearing of the lysate (30-60 min). The obtained lysates were diluted by adding 350 μL of TES and 16 μL of 5M NaCl and gently extracted twice with equal volume of 25:24:1 phenol:chloroform:isoamyl alcohol mix with room temperature centrifugation at 8,000 g for 5 min. This was followed by a similarly done extraction by an equal volume of chloroform. Finally, aqueous phase was gently mixed with 2 volumes of 96% ethanol and DNA was pelleted by centrifugation at 8,000 g, 4°C for 15 min. Pellets were rinsed with 1 mL of 70% ethanol, air dried and resuspended in 50 μL TE (10 mM Tris-HCl, 0.1 mM disodium EDTA, pH 8.0) by overnight incubation at 4°C.

Long-read sequencing libraries (SRR13062400, SRR13062401, SRR13062402, SRR13062326, SRR13062327) were prepared using Oxford Nanopore Rapid Barcoding Kit (SQK-RBK004) and pooled. The pooled library was then loaded onto R9 version flowcell (FLO-MIN106D) and sequenced in a MinION sequencer (Oxford Nanopore) for 48 hours. Sequencing data was then basecalled, quality-filtered and de-multiplexed using Guppy v3.1.5. This yielded a total of 299,317 reads of 1,882±4,697 nt length (median±IQR) for APC919/174, 27,348 reads of 11,268±24,836 nt length for 8T, 57,259 reads of 10,529±21,955 nt length for 8W, 31,238 reads of 13,673±32,494 nt length for Phr5, and 6,547 reads of 4,207±16,442 nt length for Phr5.

Trimmed and filtered Illumina reads and raw Oxford Nanopore reads were used for hybrid assemblies of the genomes of strains APC919/174, 8T and 8W using SPAdes assembler v3.13.0 in ‘careful’ mode. This resulted in complete circular contigs of 5,785,761 bp, 5,785,768 bp and 5,785,766 bp respectively. Genomes were annotated using RASTtk and PGAP pipelines and deposited in NCBI Genbank under accessions numbers CP041379, CP064941 and CP064940. Completed genomes were aligned using progressive MAUVE algorithm (version 2015-02-13) to identify regions of major recombination (**Fig. S1**).

For detection of PVR promoter inversions and variant analysis in the *in vitro* co-culture and mouse colonisation experiments (**Figs. 3d-e, S3b**) paired Illumina reads were mapped to the assembled APC919/174 as a reference. Reads were aligned using bowtie2 v2.3.4.1 in the end-to-end mode. Alignments were sorted, indexed and converted into counts tables using samtools v1.7.

For detection of genomic recombination on a single read level, sufficiently long Oxford Nanopore reads (> 1000 nt) were aligned against APC919/174 reference genome using BLASTn v2.10.0 with e-value cut-off of 1e-20. Individual alignments were retained only if length was >200 nt with identity of >90%. Next, reads were identified containing internally inverted alignments or blocks of aligned sequence corresponding to coordinates in the reference genome shifted >200 nt relative to other aligned blocks inside the same read. Such blocks of inconsistently aligned sequence were deemed as evidence of recombination events dynamically unfolding inside an individual clonal culture. Starting coordinates of these misaligned blocks were then plotted against the APC919/174 reference genome in a histogram with 1000 nt long bins and hotsposts of recombination were identified (**Fig. S2**).

### RNAseq procedures

Transcriptomic analysis of ΦcrAss001/*B. intestinalis* co-cultures was conducted for *in vitro* phage persistence experiment (**Fig. 3a, 3c**; BioSamples SAMN16809412-SAMN16809423) and one-step growth experiment (**Fig. 4a-b**; BioSamples SAMN16810643-SAMN16810654).

One millilitre culture aliquots were pelleted by centrifugation for 3 min at 17,000 g, anaerobically. Pellets were immediately lysed using 1 mL of TRIzol reagent (ThermoFisher Scientific). Total bacterial RNA was extracted using the standard manufacturers protocol. Extracted RNA (20 μL) was treated with 2U TURBO DNase (ThermoFisher Scientific), further purified using RNeasy Mini Kit (Qiagen) and assayed for RNA integrity and quantity on Agilent Bioanalyzer using 6000 RNA Nano Kit.

Stranded RNA-seq libraries were prepared from 1 μg total RNA using ScriptSeq v2 Complete kit for bacteria (Epicentre) and sequenced on Illumina NovaSeq 6000 platform at GENEWIZ. Read quality checks, filtration and trimming was performed as described above in *Shotgun sequencing and analysis of bacterial genomes*.

Reads were then mapped to ΦcrAss001 and *B. intestinalis* APC919/174 reference genomes (NCBI accession numbers MH675552 and CP041379 respectively) using bowtie2 v2.3.4.1 in the end-to-end mode. Alignments were sorted by DNA strand and indexed using samtools v1.7. Samtools ‘mpileup’ command was used to calculate read coverage per nucleotide position in both genomes. HTSeq framework v0.6.1.p1 was used to calculate aligned read counts per feature with default parameters. Read counts and coverage tables were then imported into R environment using data.table library v1.12.8 and compared between time points using DESeq2, using Wald test and parametric fit.

### Quantification and statistical analysis

Bacterial colonisation levels *in vivo* in the presence or absence of phage (**Fig. 1b**) were compared using Kruskal-Wallis test with FDR correction and were all found to be insignificantly different, except for days 31 and 38 (p < 0.05). Adsorption of crAss001 to sensitive and resistant bacterial variants (**Fig. 1b**) was compared using Kruskal-Wallis test with Bonferroni correction and was found significantly different (p < 0.05). Analysis of differentially abundant transcripts (**Figs. 3a, 4b**) was performed using DESeq2. Only genes showing significant change (p < 0.001 for host genes in in vitro persistence experiment, **Fig. 3a**, and p < 0.05 for both phage and host genes in one step growth experiment, **Fig. 4b**) between any of the consecutive time points are displayed. Bacterial genes differentially affected by mutations between the two groups of animals (phage and no phage, **Fig. S3b**) in mouse colonisation experiment were selected using exact Fisher’s test (p < 0.05 with FDR correction).

