## Supplemental Information for "Long-term persistence of crAss-like phage crAss001 is associated with phase variation in *Bacteroides intestinalis*"

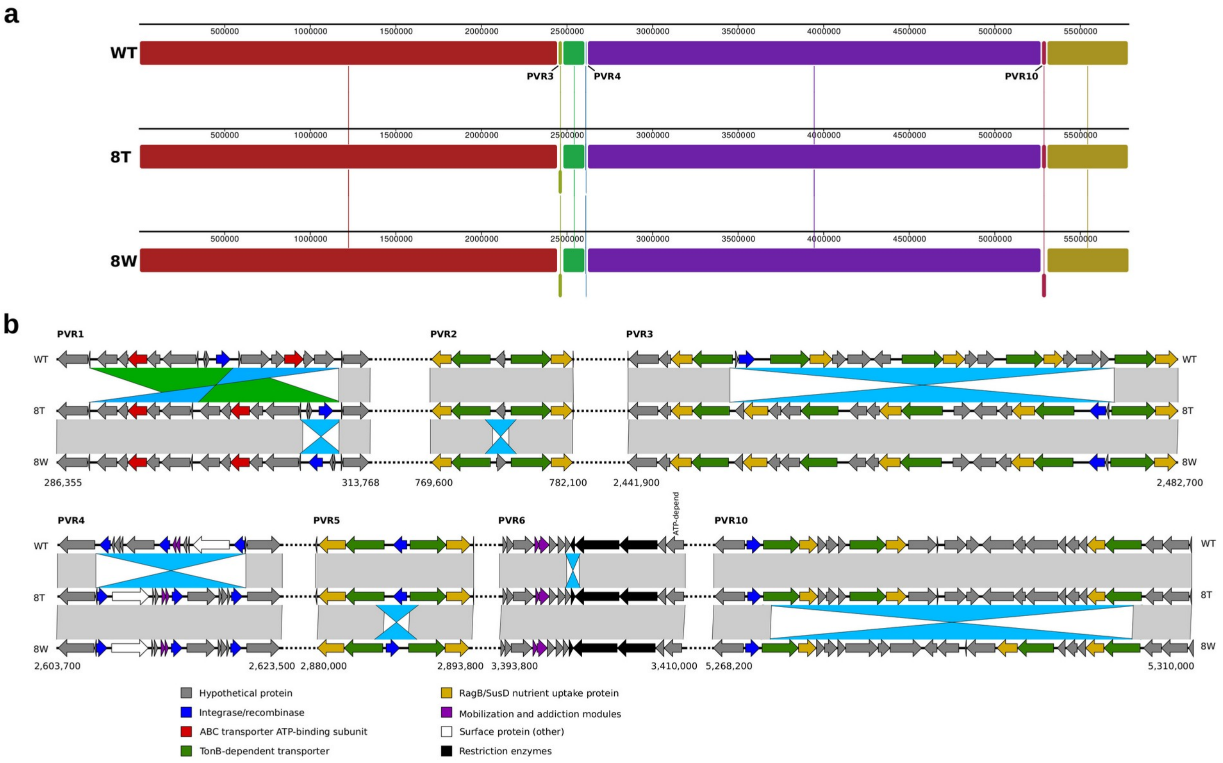


**Figure S1. Phase variation associated with major rearrangements in *B. intestinalis* APC919/174 genome. a**, MAUVE alignment of original wild-type strain, phage resistant clone 8T and spontaneous revertant clone 8W complete circular genomes showing major inversions; b, phase variable regions (PVR1-6, 10) with large inversions and translocations; protein sequence homologies (tBLASTx) between gene products are shown as coloured parallelograms.

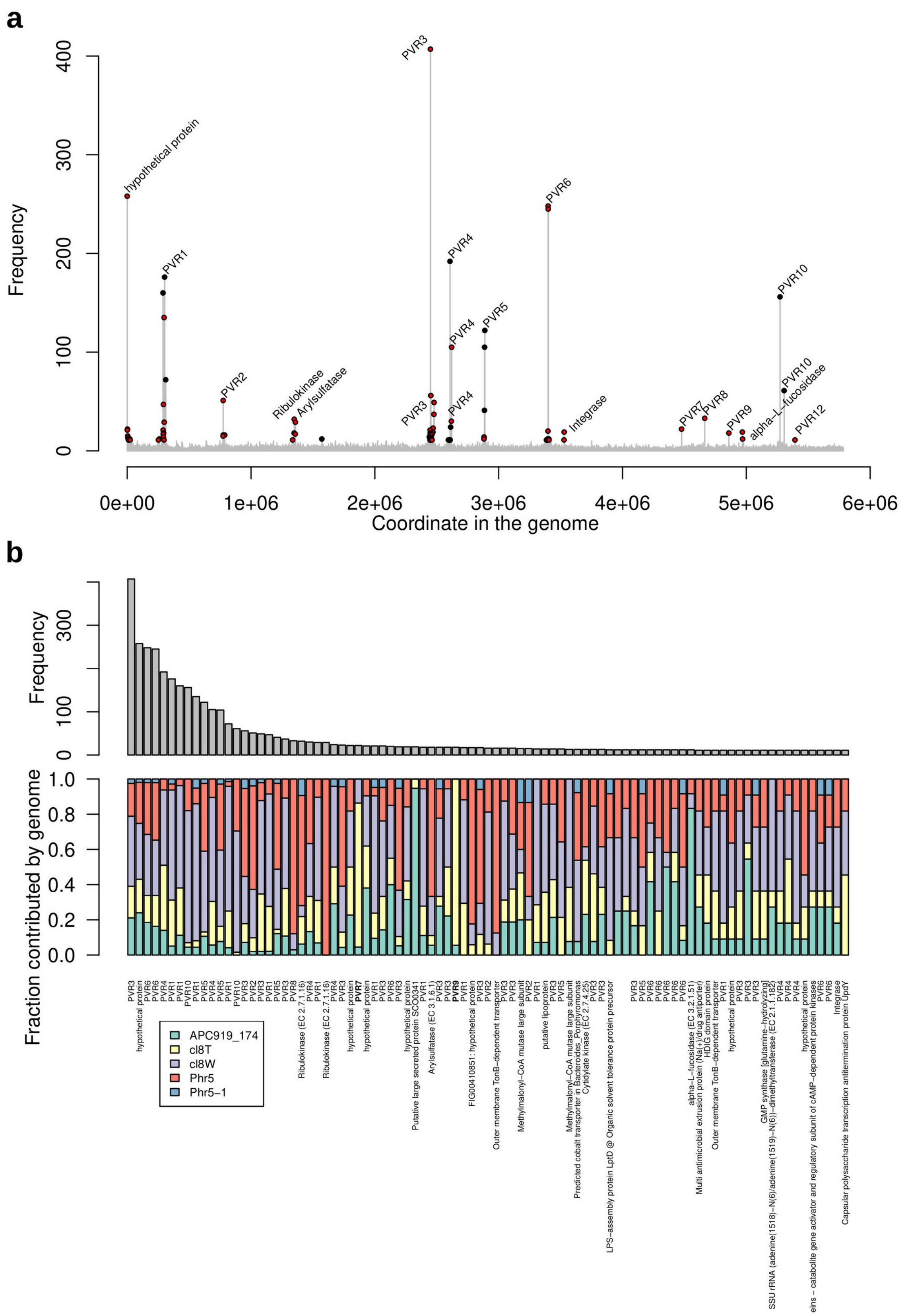


**Figure S2. Discovery of additional phase variable regions n *B. intestinalis* APC919/174 genome using Oxford Nanopore MinION long read sequencing platform. a**, Frequency of detected recombinations at a single read level (reads of at least 1000nt, with individual alignments of >90% identity and >200nt length; all inversions or shifts in coordinates >200 nt were deemed as recombinations) versus coordinates in the chromosome scaffold (histogram bin size = 1000bp); reads were pooled from sequencing of the following strains: APC919/174 WT, cl8T (phage resistant derivative), cl8W (spontaneous phage-sensitive revertant clone), Phr5 (phage resistant derivative), Phr5-1 (spontaneous phage-sensitive revertant clone); recombination hotspots were identified when >10 reads with inconsistent alignment were present per 1000bp bin; PVR regions or gene products overlapping with hotspots are marked on the plot; **b**, recombination hotspot re-assorted by frequency (top bar plot) with fraction of recombined reads originating from each of the strains (bottom stacked bar plot).

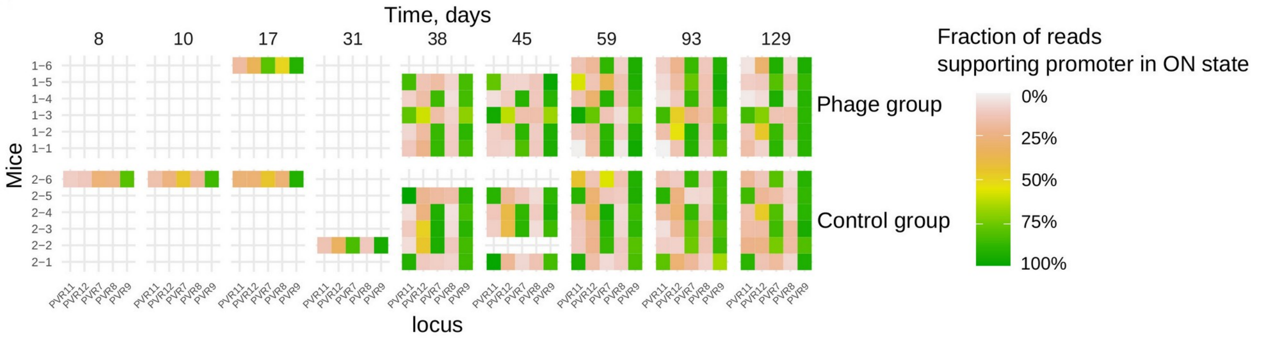


**Figure S3. Phase variation in *Bacteroides intestinalis* APC919/174 CPS operon expression in mouse colonisation experiment in the presence or absence of phage crAss001.** Displayed is the fraction of Illumina concordantly-aligned read pairs supporting orientation of invertible promoters in either ON or OFF directions; green corresponds to promoter being fully ON, red – fully OFF.

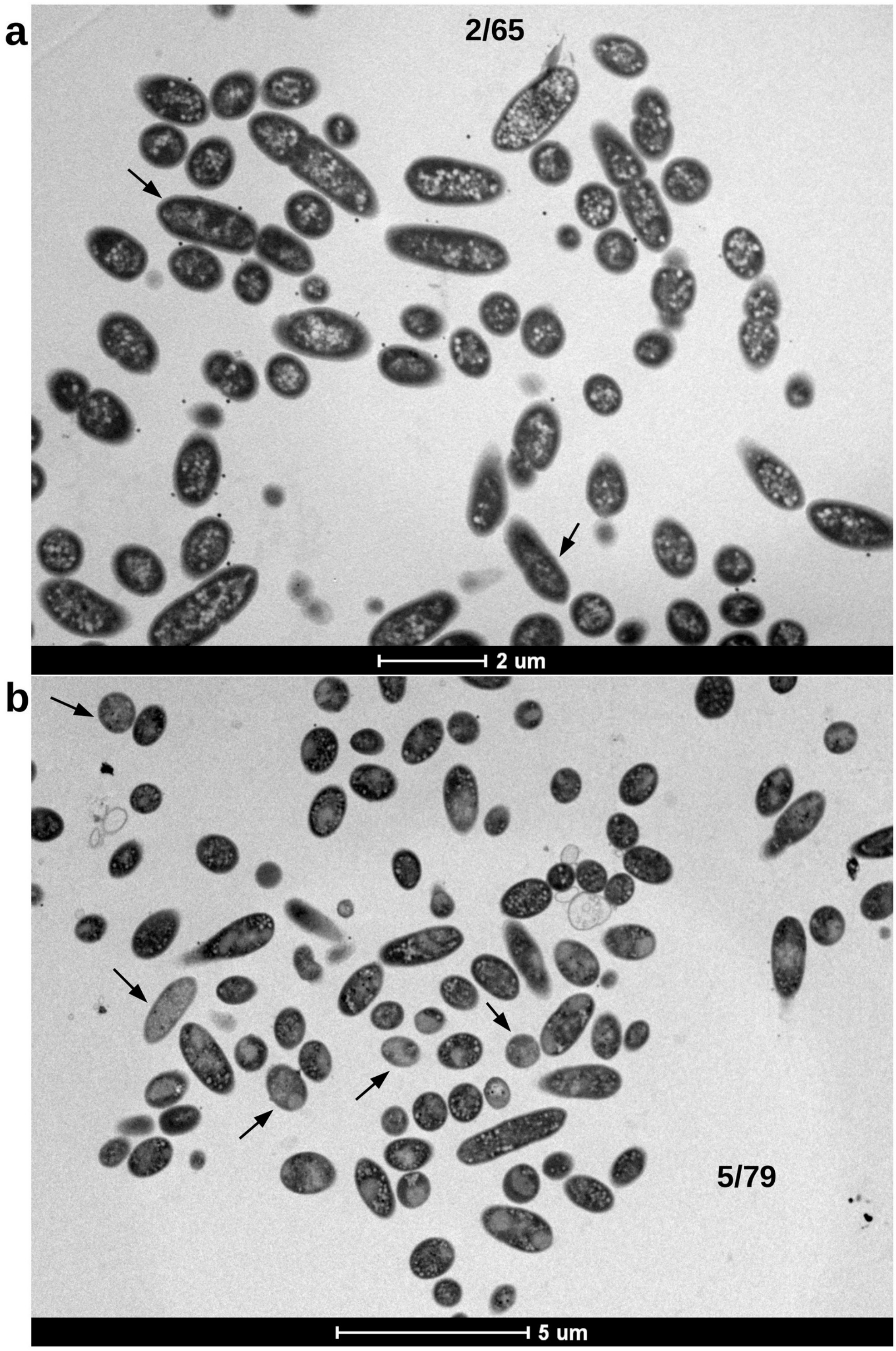


**Figure S4. TEM of *B. intestinalis* cultures infected with crAss001 at an MOI=1 reveal that >90% of cells show signs of early stages of virion assembly process 40 (a) and 90 (b) minutes after infection** (2/65 and 5/79 respectively, 6,000x and 4,200x).

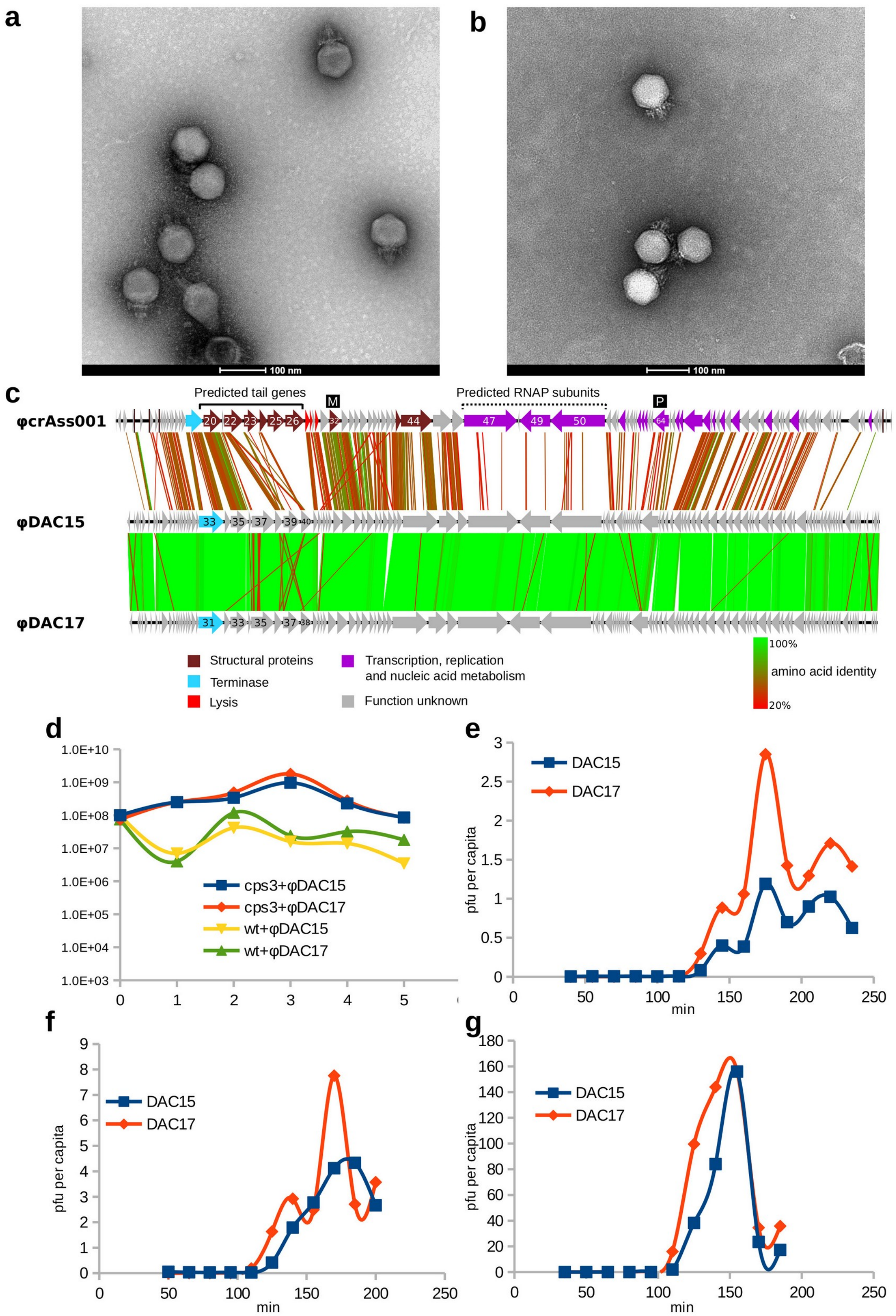


**Figure S5. Morphology, genome structure and biological properties of *B. thetatiotaomicron* phages DAC15 and DAC17 in comparison with ΦcrAss001. a**, TEM of negatively contrasted CsCl gradient-purified ΦcrAss001 particles (x62,000); **b**, TEM of negatively contrasted CsCl gradient-purified DAC15 particles (x49,000); **c**, genome comparison of ΦcrAss001, DAC15 and DAC17 highlighting overall synteny and protein sequence conservation (tBLASTx), protein sequence homologies are shown as coloured parallelograms; **d**, five-day persistence experiment of DAC15/DAC17 inoculated into early-log phase cultures of either *B. thetatiotaomicron* VPI-5482 wild-type (wt) or single-CPS3 expressing mutant (*cps3*$^{+}$), at an MOI=1; **e** and **f**, one-step growth curves of DAC15 and DAC17 in VPI-5482 wt and VPI-5482 *cps3*$^{+}$, respectively, after infection of early-log-phase cells at an MOI=1; **g**, one-step growth curves of DAC15 and DAC17 in VPI-5482 cps3 after infection of early-log-phase cells at an MOI=10.
